# Isolation of SARS-CoV-2 B.1.1.28.2 P2 variant and pathogenicity comparison with D614G variant in hamster model

**DOI:** 10.1101/2021.05.24.445424

**Authors:** Pragya Yadav, Sreelekshmy Mohandas, Prasad Sarkale, Dimpal Nyayanit, Anita Shete, Rima Sahay, Varsha Potdar, Shreekant Baradkar, Nivedita Gupta, Gajanan Sapkal, Priya Abraham, Samiran Panda, Balram Bhargava

**Affiliations:** Indian Council of Medical Research-National Institute of Virology, Pune, Maharashtra, India, Pin-411021; Indian Council of Medical Research, V. Ramalingaswami Bhawan, P.O. Box No. 4911, Ansari Nagar, New Delhi, India Pin-110029

**Keywords:** SARS-CoV-2, B.1.1.28.2, P2 variant, India, pathogenicity

## Abstract

**Background:** Considering the potential threat from emerging SARS-CoV-2 variants and the rising COVID-19 cases, SARS-CoV-2 genomic surveillance is ongoing in India. We report herewith the isolation of the P.2 variant (B.1.1.28.2) from international travelers and further its pathogenicity evaluation and comparison with D614G variant (B.1) in hamster model.

**Methods:** Virus isolation was performed in Vero CCL81 cells and genomic characterization by next generation sequencing. The pathogenicity of the isolate was assessed in Syrian hamster model and compared with B.1 variant.

**Results:** B.1.1.28.2 variant was isolated from nasal/throat swabs of international travelers returned to India from United Kingdom and Brazil. The B.1.1.28.2 variant induced body weight loss, viral replication in the respiratory tract, lung lesions and caused severe lung pathology in infected Syrian hamster model in comparison, with B.1 variant infected hamsters. The sera from B.1.1.28.2 infected hamsters efficiently neutralized the D614G variant virus whereas 6-fold reduction in the neutralization was seen in case of D614G variant infected hamsters’ sera with the B.1.1.28.2 variant.

**Conclusions:** B.1.1.28.2 lineage variant could be successfully isolated and characterization could be performed. Pathogenicity of the isolate was demonstrated in Syrian hamster model and in comparison, with B.1 variant was found more pathogenic. The findings of increased disease severity and neutralization reduction is of great concern and point towards the need for screening the vaccines for efficacy.

## 1. Introduction

Over the year of COVID-19 pandemic, Severe Acute Respiratory Syndrome Corona virus-2 (SARS-CoV-2) has accumulated severe mutations leading to the emergence of new variants. The first SARS-CoV-2 variant of concern, B.1.1.7, 20I/501Y.V1, VOC202012/01 was identified in late December in the United Kingdom (UK) which is now reported in more than 62 countries in Europe, Asia, United States of America, and elsewhere. This variant has about 17 mutations, including N501Y, P681H, 69–70 deletion; the ORF8 Q27stop mutation outside the spike protein and is adapted to be more transmissible [1]. Subsequently, another variant B.1.351, 20C/501Y.V2 was identified in South Africa with 21 mutations, including N501Y, E484K, and K417N on the spike protein, and ORF1b deletion outside the spike protein. It has been reported in Africa, Europe, Asia, and Australia with studies reporting potential immune escape [2].

During January 2021, lineage P.1, also known as 20J/501Y.V3, Variant of Concern with 17 amino acid changes which includes N501Y, E484K, and K417N on the spike protein; ORF1b deletion outside the spike protein was determined in the travelers from Brazil at Japan. The virus has been found to be widely distributed in Amazonas state of Brazil and also noticed in the Faroe Islands, South Korea, and USA [3,4]. The virulence, transmissibility and immune invasion potential of this variant remains unknown. Still, there is a concern that this variant might have facilitated the re-infections in Manus city of Amazonas which had achieved herd immunity in October 2020. Furthermore, Brazil reported another variant P.2 lineage, which has E484K mutation but not the N501Y and K417N amino acid changes in the spike protein [3].

Public health experts across the globe are rigorously following the three most rampantly spreading SARS-CoV-2 variants identified in the United Kingdom, South Africa and Brazil with concern. The mutations in receptor binding domain (RBD) of the spike protein enabled these variants to have strong affinity and binding capacity to receptor Angiotensin converting enzyme 2 (ACE2) leading to higher transmissibility [1–4].Also, many of these variants are reported to be linked to the immune escape and neutralization reduction threatening the vaccine policies and potential monoclonal antibody treatments [5,6]. Pathogenicity of the SARS-CoV-2 virus variants can be assessed in animal models. For SARS CoV-2 multiple animal models are available like ferrets, Syrian hamsters, non-human primates etc[7]. Among this, Syrian hamster is a widely used model for SARS CoV-2 which develops pneumonia and body weight loss following infection(Mohandas et al., 2020).Syrian hamsters have been used to assess the pathogenicity of many SARS-CoV-2 variants like B.1, B.1.1.7, B.1.617 [8,9].

In India, many cases of UK variants have been already reported [10]. More recently, four cases of the SARS-CoV-2 variant B.1.351 from South Africa and one case of the Brazil variant P2 lineage have been detected in the country[11]. The recent emergence of VOC B.1.617.1, VOC B.1.617.2 and VOC B.1.617.3 has caused further situation worrisome in India[12]. Increase in the COVID-19 cases has been observed during the early 2021. Considering the potential threat from emerging variants, the Government of India has continued the SARS-CoV-2 genomic surveillance of the International travelers and their contact tracing. Here, we report isolation and characterization of the Brazil P.2 (B.1.1.28.2) lineage from clinical specimens of international travelers and its impact on hamster while infected and compared with B.1 variant with known mutation of D614G.

## 2. Methods

### 2.1 Clinical specimens

The throat/nasal swabs of two COVID-19 asymptomatic cases (aged 69 and 26 years) were used for the isolation. The samples were of international travelers who returned from UK (in December 2020) and Brazil (in January 2021) respectively to India. Both the cases did not have any co-morbid conditions and were asymptomatic throughout the course of infection till recovery.

### 2.2 Cells and virus

Vero CCL81 cells grown in Minimum Essential Media (MEM) supplemented with 2% fetal bovine serum (FBS) were used. SARS-CoV-2 B.1 variant (D614G variant), NIV-2020-770 isolated from a patient’s throat/ nasal swab sample with a titre of 10^6.5^ tissue culture infective dose 50 (TCID50)/ml was used for the pathogenicity comparison study with the B.1.1.28.2 isolate [13].

### 2.3 Virus isolation and titration

Hundred microliters of the tissue homogenate/ swab specimens were added onto a 24-well plate of VeroCCL81 cells and incubated at 37°C for one hour. After washing with phosphate buffered saline and removal of media, the plates were incubated with maintenance media in a CO2 incubator at 37°C. The plates were observed daily for any cytopathic effects (CPE) using an inverted microscope (Nikon, Eclipse Ti, Japan). On observation of CPE, the supernatant was harvested and confirmed by real-time RT-PCR. Titration was performed by Reed and Muench method and titer was expressed as TCID50/ml [14].

### 2.4 Next Generation Sequencing

Genomic characterization of clinical samples and virus isolates were carried out using Next-Generation Sequencing (NGS) as per earlier explained protocols [15]. Briefly, ribosomal RNA depletion was carried on the extracted RNA using Nebnext rRNA depletion kit (Human/mouse/rat) (New England BioLabs. In, USA). Subsequently, cDNA was synthesized with the first strand and second synthesis kit. The RNA libraries were prepared using TruSeqStranded total RNA library preparation kit (Illumina, USA). The amplified RNA libraries were quantified and loaded on the MiSeq Illumina sequencing platform after normalization [6]. The genomic sequences for the clinical specimens and the virus isolates were retrieved using the reference-based mapping with the SARS-CoV-2 reference sequence (Accession Number: NC_045512.2) in CLC Genomics Workbench v20.0.4 and submitted to the public repository i.e., GISAID. The retrieved sequences were aligned with other representative SARSCoV-2 sequences downloaded from the GISAID database and a phylogenetic tree was generated with representative sequences from GISAID. General Time Reversible model with Gamma distribution of 0.05 and a boot strap replication of 1000 cycles was used to generate a Maximum-Likelihood tree.

### 2.5 Ethics statement

All the animal experiments were performed with the approval of Institutional Animal Ethics Committee and as per the guidelines of CPCSEA, India.

### 2.6 Pathogenicity study in Syrian hamsters

To understand the pathogenicity of the isolate, we intranasally inoculated 10^4^ TCID50 virus dose of Brazil P.2 (B.1.1.28.2) in 9 Syrian hamsters and observed them for 7 days for any disease and three hamsters from each group were sacrificed on day 3, 5 and 7 to study the organ viral load, antibody response and lung pathology. Simultaneously the pathogenicity of this variant was compared with a widely circulating characterized B.1 lineage SARS-CoV-2 variant in Syrian hamsters. For this, 9 hamsters were intranasally infected with 10^4^ TCID50 virus dose of B.1 lineage variant and observed for any clinical signs and studied the organ viral load, antibody response and lung pathology in sacrificed animals as defined for B.1.1.28.2.

### 2.7 SARS-CoV-2 E gene real time RT-PCR to detect gRNA and sgRNA

Nasal wash, throat swab and organ tissue samples were collected from hamsters at necropsy. Organ samples were homogenized in 1 ml cell culture media using tissue homogenizer (Eppendorf, Germany). MagMAX™ Viral/Pathogen Nucleic Acid Isolation Kit was used for RNA extraction as per the manufacturer’s instructions. Real-time RT-PCR was performed for SARS-CoV-2 E gene (both genomic and subgenomic RNA) as described earlier [16,17].

### 2.8 Enzyme-linked Immunosorbent Assay

The hamster serum samples collected on day 3, 5, 7 post virus inoculation were tested for IgG antibodies by hamster anti-SARS-CoV-2 IgG ELISA as described earlier [18].

### 2.9 Plaque Reduction Neutralization test (PRNT50)

PRNT50 was performed on serum samples of hamsters against SARS-CoV-2 B.1 variant and B.1.1.28.2 variant as per earlier described methodology [19].

### 2.10 Histopathology

Lungs samples fixed in 10% neutral buffered formalin were processed using an automated tissue processor and were embedded in paraffin. The tissues were sectioned using an automated microtome (Leica, Germany) and stained by routine hematoxylin and eosin staining. The lesions were graded as severe (+4), moderately severe (+3), minimal (+2), mild (+1) and no lesions (0) after scoring for the vascular changes, alveolar damage and inflammatory changes.

### 2.11 Data analysis

Graph pad Prism version 8.4.3 software was used for the analysis. Non parametric Mann Whitney tests were used and the p-values less than 0.05 were considered to be statistically significant.

## 3. Results

### 3.1 B.1.1.28.2 isolation and characterization

Clinical specimens inoculated in to Vero CCL-81 cells showed a typical rounding and detachment of the infected cells on 4th post-inoculation day (PID) (Figure 1). The progressive infectivity was observed with fusion of the infected cells with neighboring cells leading to the generation of large mass of cells. The presence of the replication competent virus was confirmed by Real time RT-PCR that demonstrated higher viral load in the cell culture medium on PID-3 than inoculated specimens. The virus isolate titrated at passage 2 and passage 3 demonstrated a virus titer of 10^4.5^ and 10^5.13^ TCID50/ml respectively. On phylogenetic analysis, the clinical specimens and virus isolates were found to cluster with the Brazil P2 lineage sequences. The clinical and the isolate sequences had amino acid mutations similar to the ones reported in the Brazilian P2 variant (Figure 1).

**Figure 1:**
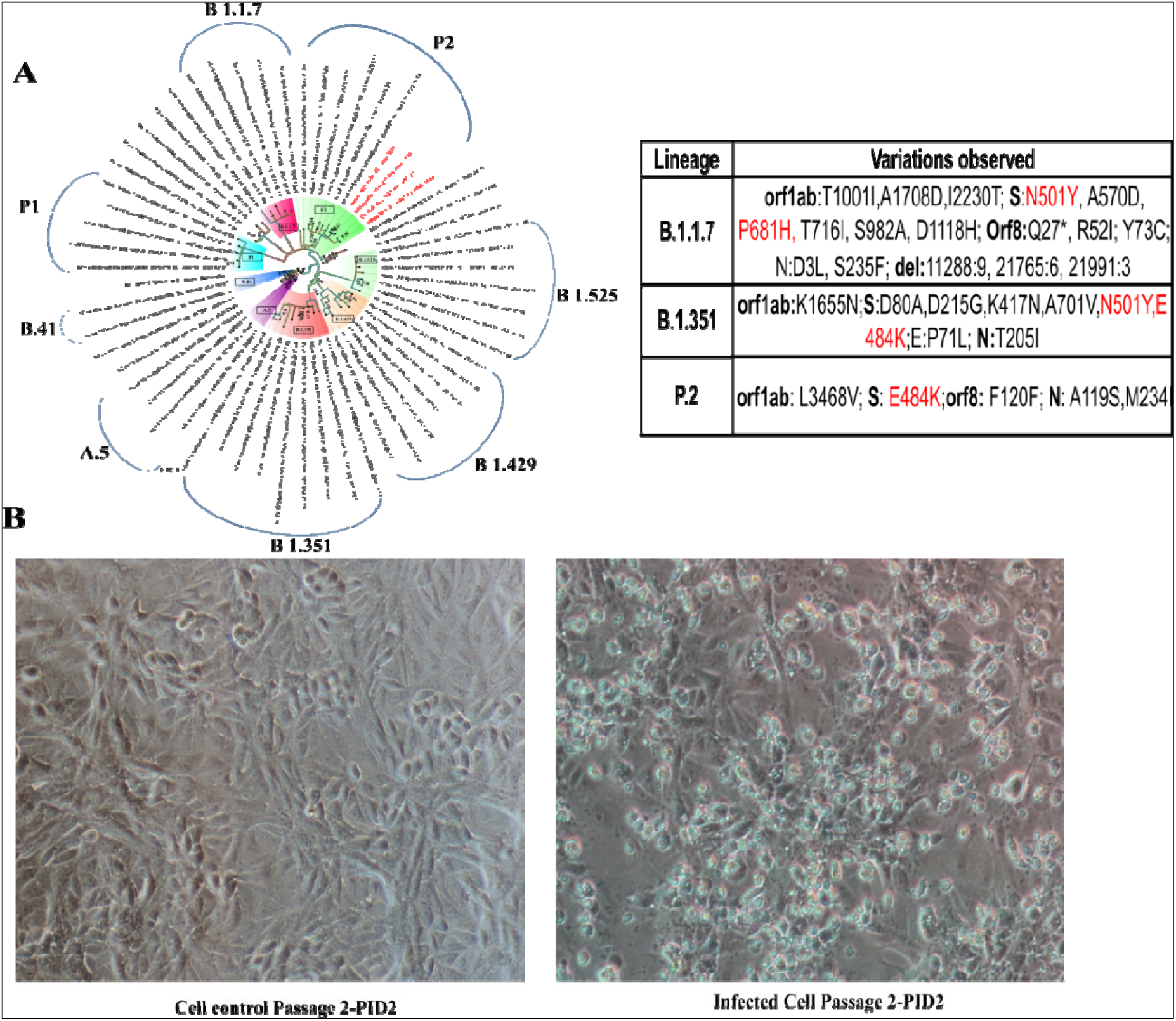
Isolation and characterization of B.1.128 P2 Brazil variants: (A) Maximum likelihood tree of the clinical samples and its isolate from the international travelers. The tree was generated using general time reversible model with a bootstrap replication of 1000 cycles. The sequences retrieved in from clinical specimens and its isolates are highlighted in red (B) The cytopathic effect observed in the Vero CCL-81 cell culture at passage-2 PID-4 compared to the cell control.

### 3.2 Pathogenicity of B.1.1.28.2 in hamsters

We inoculated 9 Syrian hamsters intranasally with 10^4^ TCID50 of the B.1.1.28.2 isolate.Body weight loss was observed in hamsters post inoculation and the average percent weight change in hamsters were −9%, −15% and −20% on day 3, 5 and 7 respectively (Figure 2A). Viral genomic RNA load was highest on day 3 and among the organs, lungs (mean = 2.7 × 10^11^ copies/ml) showed the maximum viral RNA load followed by trachea (mean= 5.9 × 10^8^ copies/ml and nasal turbinate’s(mean = 3.2 × 10^9^ copies/ml).Nasal wash and throat swab showed mean gRNA load of 1.6 × 10^9^and 3.3 × 10^8^copies/ml (Figure 2B-F). Viral gRNA load showed reduction in the further days to reach 2.4 × 10^8^ in lungs, 5.6 × 10^8^ in nasal turbinates and 2.5 × 10^8^ copies/ ml in nasal wash.

**Figure 2:**
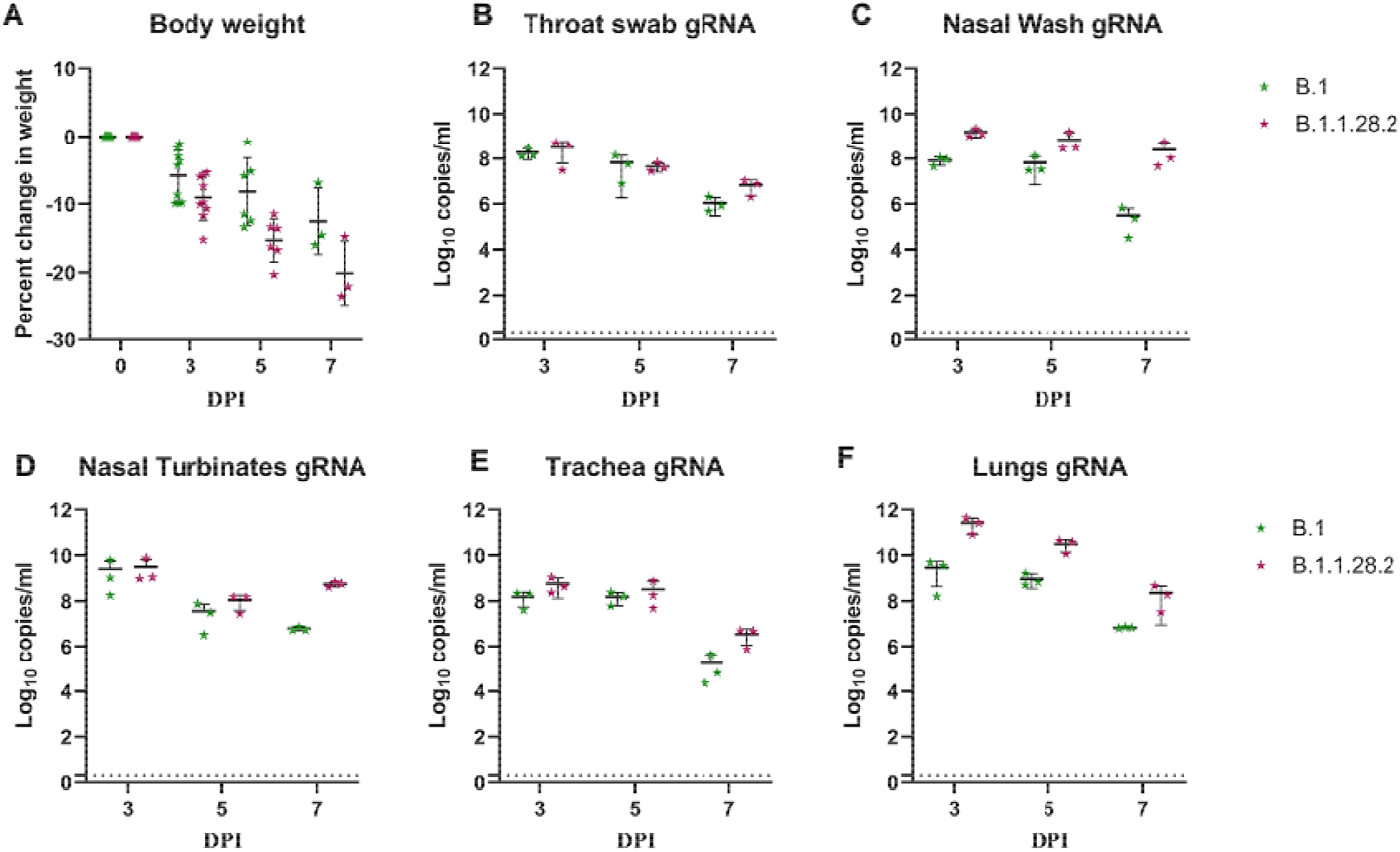
Body weight change and SARS-CoV-2 viral genomic RNA load in hamsters post inoculation by B.1 and B.1.1.28.2 lineage variants. A) Percent body weight change in hamsters post virus inoculation. Viral load in B) throat swab C) nasal wash D) nasal turbinates E) trache and F) lungs post inoculation.

Sub genomic RNA could be detected in the nasal turbinate and nasal wash samples till day 7 whereas in lungs it could be detected till day 5 in all hamsters and in only one hamster on day 7 (Figure 3). The average virus titerof 10^4.7^, 10^4.16^ and 10^2.16^TCID50/ml could be observed in lungs on day 3, 5 and 7 respectively, whereas nasal turbinate’s showed 100-fold lessertiterthan lungs on day 3 and negligible or no titer in the further days (Figure 4).

**Figure 3:**
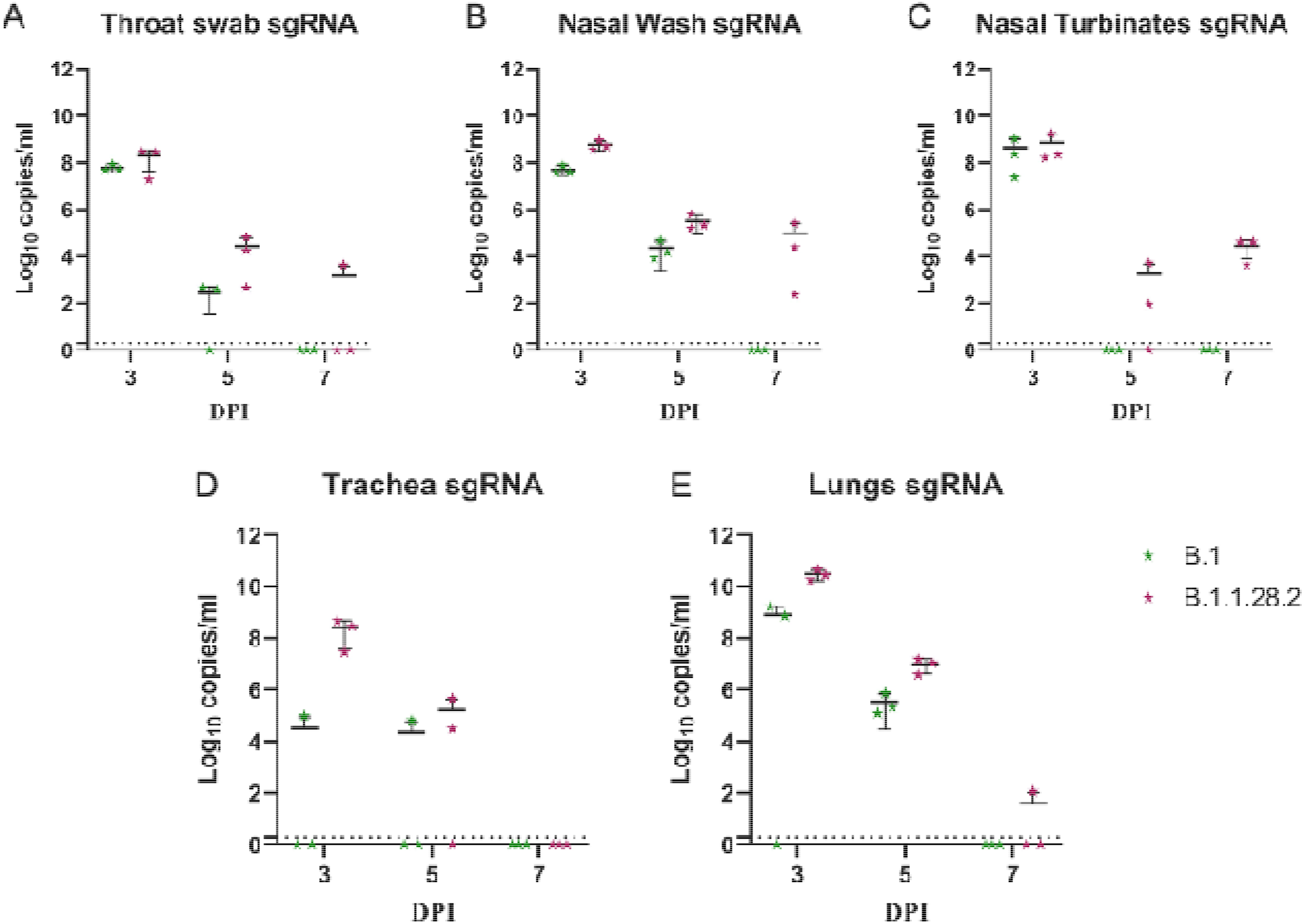
SARS-CoV-2 viral sub genomic RNA load in hamsters post inoculation by B. and B.1.1.28.2 lineage variants. Sub genomic RNA load in A) throat swab B) nasal wash C) nasal turbinates D) trachea and E) lungs post inoculation.

**Figure 4:**
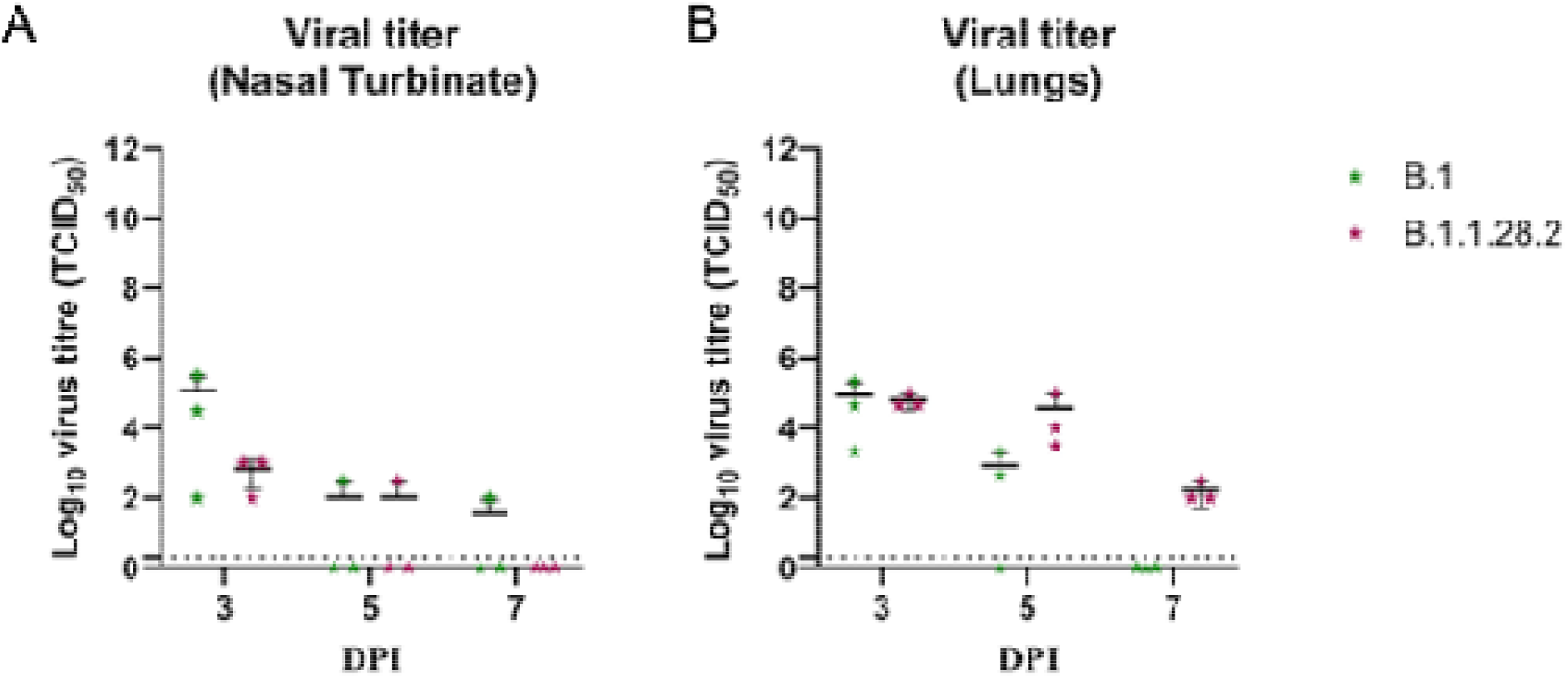
Viral load by titration. Viral load in A) nasal turbinates and B) lungs post infection by SARS-CoV-2.

Anti-SARS-CoV-2 IgG antibodies could be detected in the serum by day 7 in all hamsters by ELISA (Figure 5A).Average neutralization titer of 538.1 and 599.1 were observed with B.1 variant and B.1.1.28.2 variants respectively against B.1 variant. In case of neutralization with B.1.1.28.2 variant an average titer of 240.2 and 1256.3 were observed for B.1 and B.1.1.28.2 variants (Figure 5B, 5C). The variant induced severe lung pathological changes in lungs which includes vascular congestion, hemorrhages, interstitial septal thickening with pneumocyte hyperplasia, alveolar consolidation, perivascular, peri-bronchial and interstitial mononuclear cell infiltration. The pneumonic changes were of mild to moderate grade on day 3 which progressed to severe changes on day 7 (Figure 6A-C, Table 1).

**Figure 5:**
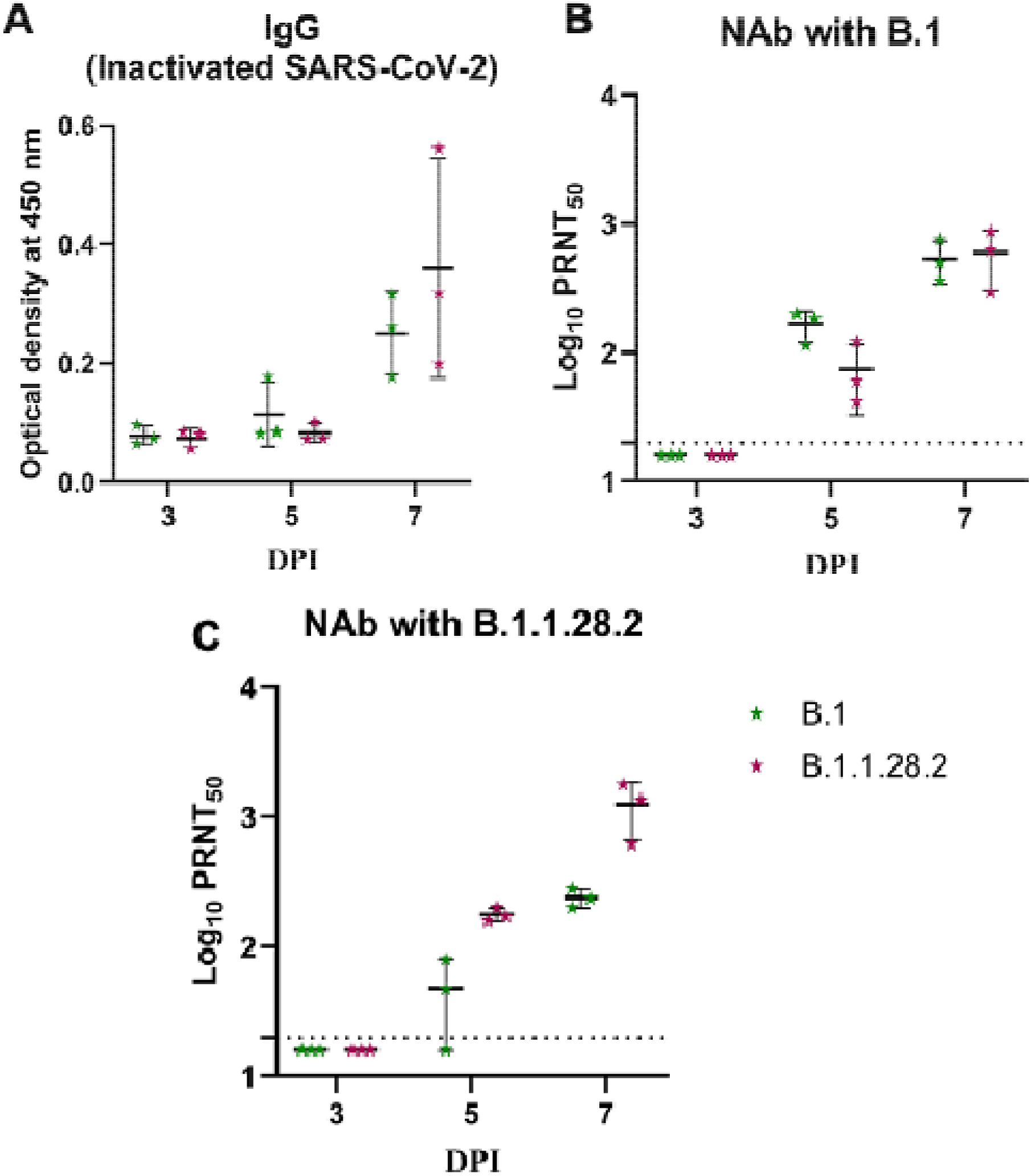
Comparison of immune response in hamsters post B.1.1.28.2 and B.1 infection. A)IgG response in hamsters measured by inactivated SARS-CoV-2 IgG ELISA. B) Neutralizing antibody response in hamsters against B.1 variant C) Neutralizing antibody response in hamsters against B.1.1.28.2 variant.

**Figure 6:**
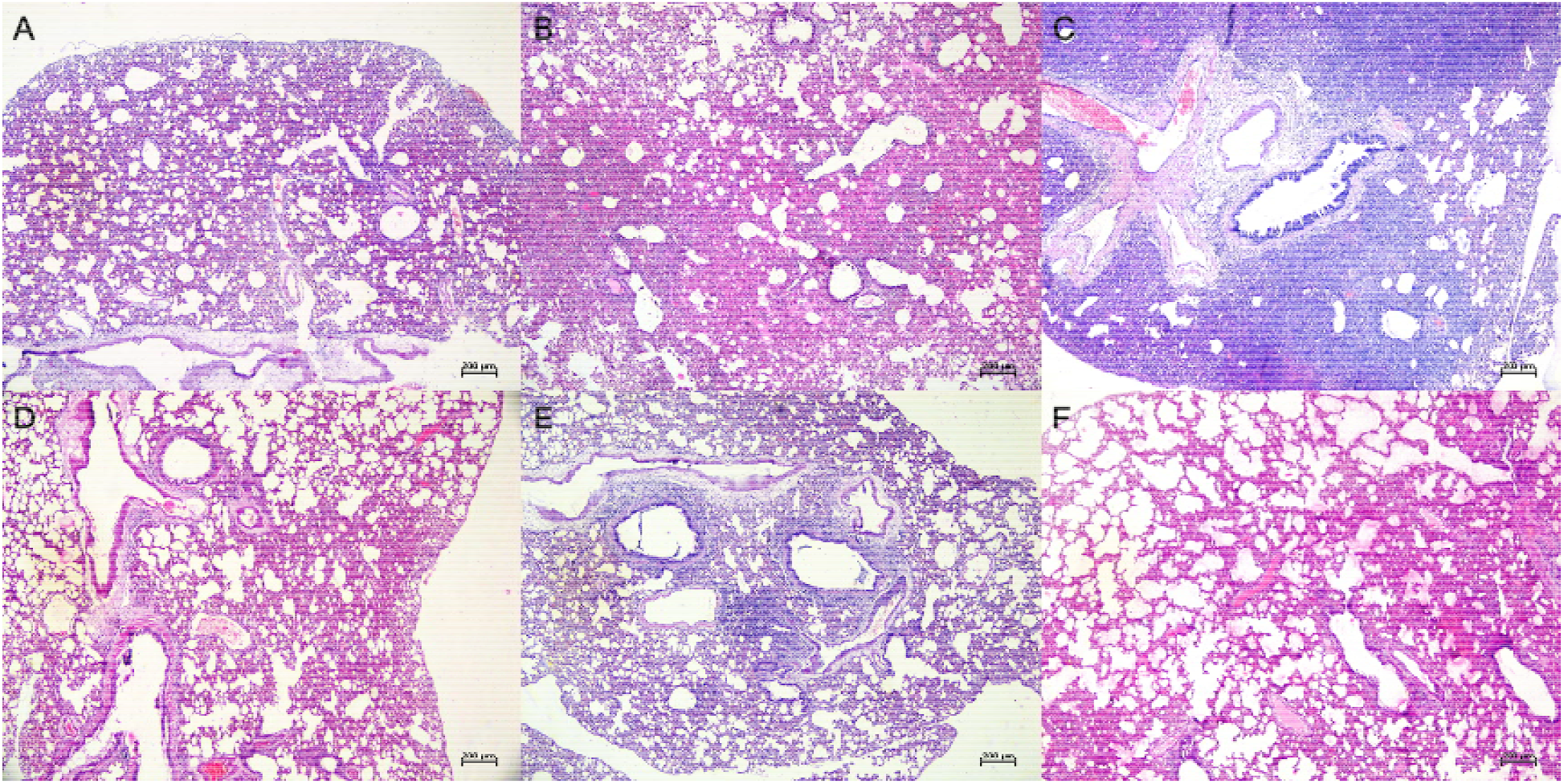
Histopathological changes in lungs post inoculation by B.1 and B.1.1.28.2 lineage variants. Lungs of hamsters infected with B.1.1.28.2 showing A) diffuse alveolar capillary congestion and alveolar septal thickening on day 3, B) diffuse capillary congestion, haemorrhages, consolidation and alveolar septal thickening on day 5 C) diffuse mononuclear infiltration in the alveolar parenchyma, congested alveolar capillaries and hemorrhages. Lungs of hamsters infected with B.1 variant showing D) engorged alveolar capillaries and of thickened alveolar septa on day 3 E) peribronchial mononuclear cell infiltration, alveolar capillary congestion and septal thickening on day 5 F) an area of consolidation with alveolar hyaline membrane formation, congestion and mononuclear cell infiltration in the alveolar septa on day 7.

### 3.3 Comparison of pathogenicity ofB.1.1.28.2 isolate with B.1 variant in infected hamsters

Although the B.1.1.28.2 lineage variant infected hamsters induced more weight loss in hamsters in comparison to the B.1 variant, it was not statistically significant (Figure 2A). A hundred-foldlesser average viral gRNA load was observed in lungs (2.9× 10^9^ copies/ml) and the nasal wash (8.3 × 10^7^copies/ ml) on day 3 in case of B.1 variant, although similar viral copies/ml were observed in nasal turbinate’s for both variants. Virus shedding through the nasal wash in B.1 variant was 8.3 × 10^7^, 6.8 × 10^7^ and 3.2 × 10^5^copies/ml on day 3, 5 and 7. On titration, lungs samples of hamster infected with B.1 variant showed an average titer of 10^4.4^ and 10^2.95^ TCID50/ml on day 3 and 5. No live virus could be detected in lungs samples of B.1 variant on day 7indicating persistent virus replication in case of B.1.1.28.2 variant where it was detected till day 7.Subgenomic RNA were detected in lungs samples only on day 3 and 5 for B.1 variant. The pneumonic changes observed in lungs induced by B.1 variant was mild in comparison (Fig6D-F, Table 1).The increased pathogenicity of the B.1.1.28.2 variant isolate as evident by the increased body weight loss, high viral load in lungs and severe lung pathology.

## 4. Discussion

The global spread of emerging SARS-CoV-2 variants, including those of the B.1.1.7(20I/501Y.V1, ‘UK variant’), B.1.351 (20H/501Y.V2, ‘South African variant’), B.1.1.28 lineages (sub-clades P.1 and P.2, ‘Brazilian variants’) and B.1.617 lineages has heightened concerns about enhanced transmissibility, virulence, and attenuation of susceptibility to humoral immunity elicited by natural infection or vaccination. Lack of clinical data and transmissibility studies of the B.1.1.28 P.1 and P.2 variants, make the course of infection in affected individual unpredictable. This also emphasizes for the specific precautions for the isolation and the management of these cases.

Here we assessed the pathogenicity of the isolate and observed disease in hamster model characterized by the weight loss and lung pneumonic changes. COVID-19 in hamsters is characterized by respiratory signs, body weight loss and pneumonia[20]. Hamster model has been used to assess the infectivity and pathogenicity of SARS-CoV-2 VOCs like B.1.1.7, B.1.351, B.1.617etc[18,21]. The variant specific differences can be evaluated with this model using criteria’s like viral load, histological scoring, inflammatory cytokine level etc[18,21]. B.1.1.28.2 variant induced body weight loss and supported viral replication in respiratory tract. Here, live infectious virus could be detected till day 7 in lung samples whereas in nasal turbinates it was detected only on day 3 and with 100-fold lesser titer indicating the virus predilection to the lower respiratory tract. Viral gRNA shed through nasal wash was higher in case of B.1.1.28. High viral shedding through nasal wash could be linked to the increased transmission efficiency speculated for this variant.

In comparison with B.1 variant, the pathogenicity of the isolate was more with severe pneumonia.Although, variants of concern are speculated to be linked to increased disease severity and transmission, *in vivo* experimental studies in case of B.1.1.7 and B.1.351 showed less or comparable disease severity with earlier SARS-CoV-2 strains[18,21]. B.1.1.7 variant showed mild lung pathology in hamster model in comparison to B.1 in our earlier study [18]. Also B.1.1.7 and B.1.351studies in hamster and non-human primate models did not show any increased disease severity[18,21,22]. In contrary, here we observed severe pneumonic changes associated with a variant of interest i.e., Brazil P2 variant.

Since the mutation in the E484 site in the RBD known to have the large effect on binding and neutralization of the SARS-CoV-2 is present in the B.1.1.28.2 variant, we assessed the neutralization potential of the hamster sera. Comparable neutralization efficiency was shown by both B.1 and B.1.1.28.2 variants infected hamster sera against B.1 variant, whereas reduction was seen in the neutralization titreof B.1 infected hamster sera against B.1.1.28.2. This finding is in line with the earlier reports of neutralization reduction in case of monoclonal antibody treatment and post vaccination sera by P.2 variant [6, 23]. The neutralization efficacy of B.1.1.28.2 variant was assessed with the convalescent sera of COVID-19 infected individual sera and BBV152 two-dose vaccinated individuals to observe about two-fold reductions in neutralizing titer against B.1.1.28.2 variant [24].

Hence genomic surveillance of SARS-CoV-2 would help in early identification of the variants of concern specifically when it is present at a low frequency. This would help in implementation of various control measures to curb the transmission of SARS-CoV-2 variants in the country. It would also allow quick assessment of their prevalence across the globe. With the research underway to determine the neutralization potential of currently available COVID-19 vaccine, we still need to continue to follow the non-pharmaceutical interventions such as use of masks, physical distancing, hand hygiene and avoiding the public gatherings which would check the transmission of these new variants to a great extent.

## 5. Conclusions

Here we report the isolation of the B.1.1.28.2 variant which was found more pathogenic in hamsters producing severe pneumonia in comparison with B.1 lineage variant. The B.1 variant infected hamster sera showed reduced neutralization against B.1.1.28.2. The study points towards the necessity of genomic surveillance and characterization of the SARS-CoV-2 variants to understand its pathogenicity and immune escape potential for preparedness.

## Acknowledgements

We gratefully acknowledge the team member of Maximum Containment Facility, ICMR-NIV, Pune including Manoj Kadam, Abhimanyu Kumar, Deepak Suryavanshi, Dr. Abhinendra Kumar, Dr. Rajlaxmi Jain, Mrs. Savita Patil, Mrs. Triparna Majumdar, Ms. Pranita Gawande, Mrs. Ashwini Waghmare, Mr. Yash, Ms. KaumudiKalele, Ms Manisha Dudhmal, Ms. Jyoti Yemul, Mr. VishwajeetDhanure and MsUjaini Shah from Influenza department for providing excellent technical support. We also acknowledge the support received from Dr. Chandrasekhar Mote, Assistant professor, Veterinary College, Shirwal, Maharashtra for the histopathology.

## Author contributions

PDY and SM contributed to study design and paper writing, PDY, NG and VP contributed to patient data collection, genomic data analysis and interpretation. SM performed animal experiments and experimental data collection. PDY, AS, DAN contributed to data analysis and interpretation, writing and critical review. RRS contributed to patient data collection, writing and critical review. GS, PS and SBcontributed to laboratory investigations. PA,SP and BB contributed to writing and critical review of the manuscript.

